# Associative Visual Memory in Aphantasia: Evidence for Intact Object and Spatial Memory, Metacognitive Awareness, but Different Strategies

**DOI:** 10.64898/2026.07.08.736656

**Authors:** Rebecca Keogh, Zoey Isherwood, Anina N. Rich

## Abstract

Many forms of memory are thought to rely on visual imagery, but individuals who report lacking visual imagery (aphantasia) can still perform various memory tasks. There is, however, evidence that aphantasia may lead to less detailed autobiographical memories, suggesting there may be deficits in the underlying cognitive processes that support personal memories. One such process is associative memory, which requires binding of different types of information. Here, we tested whether associative visual memory is intact in aphantasia. We assessed 72 self-identified individuals with aphantasia and 77 controls who reported having visual imagery. Participants completed an associative memory task which involved memorising displays where a unique object in a specific location was associated with a particular colour fixation point. Individuals with aphantasia performed equivalently to controls for object locations and outperformed controls on the associated object-identity. In addition, whereas controls were significantly worse at remembering associated object-identity than object-location, individuals with aphantasia showed no such difference. Both groups showed good metacognitive performance evidenced by a positive correlation between confidence and accuracy; there were no significant differences in confidence between the groups. Reported strategies varied between groups: a large proportion of control participants reported using visual imagery and self-reported use of imagery positively correlated with performance. Conversely, individuals with aphantasia mostly reported using nonvisual strategies to remember the associations. Overall, the findings suggest that individuals with aphantasia can form associative memories using nonvisual strategies. Thus, difficulties with autobiographical memory in aphantasia seem unlikely to be due to fundamental issues with associative memory.

## Introduction

Some people report that they are unable to visually imagine, known as aphantasia (Zeman et al., 2015). Since the term aphantasia was introduced, this phenomenon has attracted substantial interest from the public, philosophers, psychologists and cognitive scientists (Keogh, Pearson, et al., 2021). Visual imagery, like so many interesting cognitive processes, is inherently subjective, making it difficult to distinguish between differences in reported abilities that may be due to actual differences in mental imagery and differences that may reflect variation in metacognitive insight, or interpretive bias. In recognition of these known challenges, studies have investigated whether a self-reported lack of imagery is reflected in behavioural, physiological or neural measures. Individuals with aphantasia, relative to control participants who report having imagery, show reduced imagery-based priming of binocular rivalry (Keogh & Pearson, 2018, 2024; Monzel et al., 2025), attenuated skin conductance response to imagined fear inducing scenarios (Wicken et al., 2019), and diminished pupillary light responses to imagined images of varying brightness (Kay et al., 2022). Further, neuroimaging work has shown reduced functional connectivity between frontal and visual regions at rest (Milton et al., 2021) and altered connectivity between frontal regions and high-level ventral temporal cortex during mental comparison tasks (Liu et al., 2025). Recent fMRI decoding studies also suggest that while early visual areas may be recruited during attempted imagery in aphantasia, this activity does not resemble perception (Chang et al., 2025). Other studies have shown that individuals with aphantasia can perform normally on tasks commonly thought to use visual imagery, such as visual mental comparison tasks (Liu & Bartolomeo, 2023; Milton et al., 2021). Overall, although the field is still relatively young and there are ongoing debates about the neural underpinnings of aphantasia and how best to categorise it (Bartolomeo & Arcangeli, 2026; McKilliam & Kirberg, 2025; Michel et al., 2025; Zeman, 2024, 2025), converging evidence from different methodological approaches suggest that these self-reported differences in imagery ability reflect a genuine phenomenon and differences in experience.

Beyond its phenomenological interest, aphantasia has received attention because visual imagery is theorised to play an important role in a range of cognitive functions and mental health conditions (Holmes & Mathews, 2010; Pearson, 2019; Pearson et al., 2015). In particular, visual imagery has been proposed to support various types of memory, from short term memory (e.g., visual working memory: Keogh and Pearson (2011, 2014); Tong (2013)) to prospective memory (thinking about the future: Anderson et al. (2012); Schacter et al. (2008)) and autobiographical memory (recollection about one’s past: Brewer and Pani (1996); Greenberg and Knowlton (2014)). Natural variation in imagery ability, including the lack thereof, provides an opportunity to test these claims. If individuals with aphantasia are impaired on tasks that index these forms of memory, this would suggest that visual imagery is indeed *required* for them. However, a number of studies have found that individuals with aphantasia are able to perform classic short-term visual working memory tasks just as well as, or in some cases better than, individuals who report having visual imagery (Keogh, Wicken, et al., 2021; Pounder et al., 2022), although see also (Monzel et al., 2022). Additionally, mental rotation, which has been previously used to measure visual imagery ability, particularly in clinical settings, appears to be unaffected in aphantasia (Kay et al., 2024; Milton et al., 2021; Pounder et al., 2022; Zeman et al., 2010). These findings have led to two main explanations for preserved performance despite a lack of visual imagery. First, imagery might be just one strategy that can be used to perform these tasks, and other non-imagery strategies can support memory formation (Kay et al., 2024; Keogh, Wicken, et al., 2021). Second, individuals with aphantasia rely on the same representations as people with imagery, but they are unable to consciously access them (Michel et al., 2025).

Autobiographical memory, which involves recalling personal episodes from one’s own life, is one area where there is converging evidence suggesting that visual imagery may be essential. Individuals with aphantasia report having poorer autobiographical memories than their peers with imagery (Dawes et al., 2020; Zeman et al., 2015). This self-reported reduction is consistent with results from studies using the autobiographical memory interview, which show personal memories are less detailed in aphantasia compared with participants who report being able to visually imagine (Dawes et al., 2022; Milton et al., 2021). Additionally, a function magnetic resonance imaging (fMRI) study showed that when remembering past scenarios, individuals with aphantasia demonstrated reduced hippocampal activity, and different connectivity between the hippocampus and visual areas of the brain, compared with individuals who reported having visual imagery (Monzel et al., 2024). In addition, imagery ability has been found to be positively associated with the reliving component of autobiographical memory in individuals who report having visual imagery (Greenberg & Knowlton, 2014; Rubin et al., 2003). Thus, it seems that visual imagery may be necessary to support recall of personal memories. It could be that a lack of imagery reduces the sensory vividness of personal recreations, thus decreasing their potency. However, autobiographical memory is complex, drawing on semantic, associative and episodic memory. There are, therefore, multiple candidate processes that could rely on visual imagery involved in autobiographical memory leading to poorer memory for individuals with aphantasia.

Episodic memory in particular has been proposed to be affected by having aphantasia, and it has even been proposed that aphantasia should be thought of as an episodic memory disorder (Blomkvist, 2023; Blomkvist, 2025), but see Bergmann and Ortiz-Tudela (2023) for an alternate view. There is at least one study that suggests aphantasia affects other forms of episodic memory beyond personal autobiographical memories. Dando et al. (2023) had participants watch a video of a crime and found that when they were later tested on their memory of these videos, eye-witness accounts from individuals with aphantasia were less complete (i.e. they recalled less details) when they were asked to freely recall their memory of a video compared to individuals with imagery (Dando et al., 2023). However, when probed with specific questions about previously reported content, their accuracy was comparable to controls. This pattern suggests reduced spontaneous recall richness rather than impaired accuracy, and may reflect differences in encoding, retrieval or elaboration processes rather than a global deficit in episodic memory.

Episodic memory depends on the binding of event elements into integrated representations, including associations between items or people and their spatial context. Interestingly, while individuals with aphantasia report that they have no object imagery, their spatial imagery tends to be intact (Bainbridge et al., 2021; Dawes et al., 2020; Keogh & Pearson, 2018). If you have visual imagery, it might be difficult to conceive of spatial imagery existing independently from object imagery. This separation does, however, fit with the general visual hierarchy. During perception, visual information enters through the retina and is projected retinotopically onto primary visual cortex (V1). From these retinotopic representations in early visual areas, this information is passed on to more specialised temporal regions (‘ventral stream’) and parietal regions (‘dorsal stream’) (Ungerleider & Haxby, 1994). Object information is represented primarily in the temporal lobes, whereas representation of space involves the parietal lobes and dorsal regions (the ‘what’ vs ‘where’ pathway division), although of course these areas interact. It is therefore plausible that spatial and object aspects of imagery are similarly separable (Blajenkova et al., 2006). This raises the question of whether reduced object imagery in aphantasia influences the formation and/or reconstruction of associative representations underlying autobiographical memory.

Here, we aimed to test associative memory in aphantasia using an online paradigm that provides a measure of memory for object associations separately from memory for associated spatial locations. We tested participants with self-reported aphantasia and a control group reporting typical imagery. Participants were asked to remember specific images (radial frequency patterns) at defined polar angles on a computer screen, allowing us to separately measure memory for ‘what’ (object identity) and ‘where’ (spatial location), and their association with a given colour. This design allows us to examine whether a lack of visual imagery results in reduced associative memory performance, and whether this is driven by object or spatial components of the memory, providing insight into the mechanisms underlying reduced autobiographical memory in aphantasia.

We predicted that individuals who reported having aphantasia would be able to perform the task but that their performance would be poorer than control participants on the object memory component of the task only; no group differences were expected for spatial memory performance. We also hypothesised that individuals with aphantasia may be able to achieve performance levels comparable to controls by the end of the study, but this would require additional learning to form the object associations. If this is the case, we expected they would be less accurate than controls during the initial blocks of the task but not later blocks. Preliminary data from a related unpublished study suggested that individuals with aphantasia report lower confidence in their memory performance. For this reason, we also collected confidence ratings, expecting that participants with aphantasia would report lower confidence in their memories compared with control participants. We also assessed the relationship between confidence and performance, as well as reported cognitive strategies, to determine whether individuals with aphantasia had metacognitive insight into their performance.

## Methods

This study was pre-registered: https://osf.io/ahuqm

### Participants

We aimed to recruit a minimum of 100 participants with aphantasia (before exclusion criteria were applied). Participants who identified as having aphantasia were recruited through the online Australian Aphantasic Database (individuals in this database are not only from Australia) and individuals went into a draw to win a $200 AUD gift voucher for participating. 113 participants from the Australian aphantasic database participated in this study.

Non-aphantasic individuals were recruited from undergraduate students based at Macquarie University and given course credit for their participation. They formed our control group of 104 participants.

Participants were excluded for the following reasons (all pre-registered): 1. Participants with performance below 50% correct for both the object and spatial memory tasks. 2. Aphantasic participants who scored higher than 32 on the Vividness of Visual Imagery Questionnaire, in line with previous conventions (Dawes et al., 2020; Keogh & Pearson, 2024). 3. Control participants who scored below 32 on the VVIQ. 4. Any participants who reported using a strategy such as drawing the images or anything else that could be interpreted as ‘cheating’. 5. Any participant who did not move the confidence sliders or clicked the same option on every trial (i.e., selected ‘correct’ for every trial). 6. Participants who did not complete all parts of the experiment.

After these exclusions, data from 72 individuals with aphantasia and 77 control participants with visual imagery were analysed. For the cognitive strategy component of the study, data are missing for one individual with aphantasia, leaving 71 individuals with aphantasia and 77 individuals with visual imagery for this part of the analysis.

For the aphantasia group, the average age was 46.81 years (SD = 13.37, range 18-78) while the control group average age was 21.56 years (SD 7.26, range 18-60).

In aphantasia, the Vividness of Visual Imagery questionnaire (VVIQ (Marks, 1973)) has become the standard for characterising participants and there are currently no specific aphantasia screening questionnaires. Thus, although there are some issues in terms of how different participants might interpret the scale, we report the standard scores for our participants here. For the aphantasia group, the average VVIQ score was 18.28 (median = 16, SD = 3.82; range: 16-32) and the control group average was 64.07 (median = 65, SD = 10.04; range: 39-80), see supplementary materials figure S1 for the groups VVIQ data. Given our exclusion criteria, unsurprisingly the aphantasia group VVIQ was lower than the control group (W = 5544, *p* < .001, BF_10_ = 5.54×10^6^).

In addition to the VVIQ, all participants were asked *‘Do you identify as having aphantasia? Aphantasia is the inability to visualise AKA having a blind mind’s eye*.*’* and were given the following options on a 5-point Likert scale: *Definitely not, Probably not, May or may not, Probably yes*, and *Definitely yes*. In the aphantasia group, 57 participants responded ‘*Definitely yes’*, while 15 responded *‘Probably yes’*. In the control group, 1 participant responded *‘Definitely yes’* and 1 responded *‘Probably yes’* but both scored mid-range scores on the VVIQ (57 and 61) so we did not exclude them from the control group. The rest of the control participants responded with ‘*May or may not’* (N = 3), ‘*Probably not’* (N = 13) or ‘*Definitely not’* (N = 59).

### Procedure

This experiment was completed on participants’ own computers online (using Qualtrics). Participants completed demographic questions, the self-identification question regarding aphantasia, and the VVIQ, followed by an associative memory task based on (Favila et al., 2022). During the associative memory task, participants had to learn the association between the colour of a fixation dot and a specific radial pattern (object association), presented in a specific location (spatial-location association). There were two components to this task: the learning blocks and the test blocks. Participants completed 4 runs of both the learning and test blocks in an interleaved fashion (i.e., a learning block followed by test block, cycled through 4 times).

### Learning blocks

Participants had to learn and remember 4 different displays. The four displays were comprised of a coloured fixation point plus a specific radial frequency pattern at a specific polar angle (Figure 1 left panel and supplementary materials). Each display was presented for 2 seconds (no interstimulus interval) and there were 16 stimuli in total (4 unique combinations each presented 4 times), shown in a randomised order.

**Figure 1.**
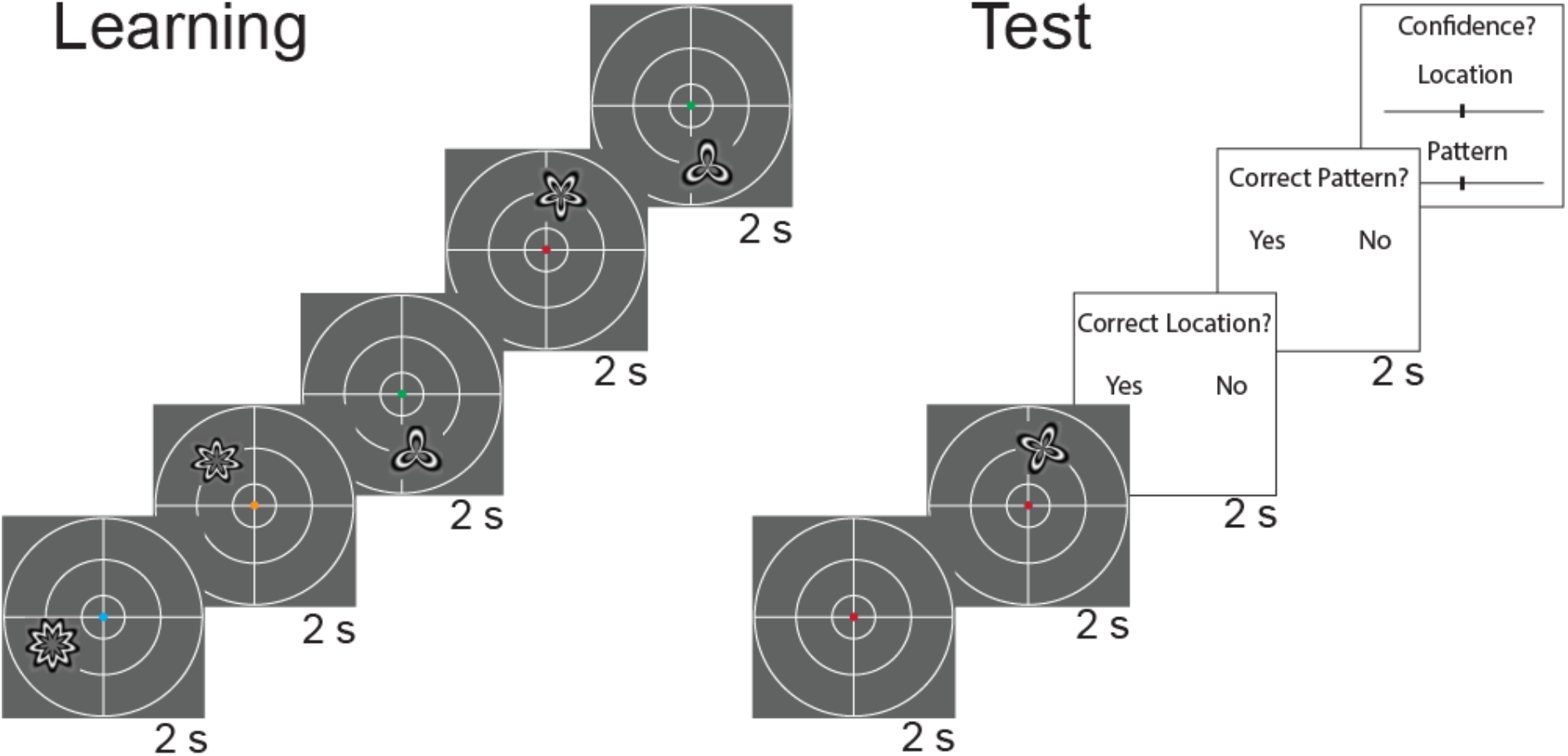
Experimental timeline. The learning task is presented on the left and the test task on the right. During the learning phase, each display was presented for 2 seconds. During the test phase participants were presented with a coloured fixation point for 2 seconds, and then a display with an additional stimulus, which could be the correct or incorrect object and location for that cue colour, for 2 seconds. They were then given 2 seconds to decide whether the location and object (pattern) was correct for the given colour and gave a rating on how confident they were with their decisions.

### Test blocks

After each learning block, participants were shown a test display (see right panel of Figure 1). There were 4 different types of test displays (see supplementary materials for all stimuli): 1. Complete match (same as the learned display); 2. Object match/spatial-location mismatch (correct radial frequency pattern associated with the central fixation colour but presented in a different location); 3. Object mismatch/spatial-location match (incorrect radial frequency pattern but presented in the correct spatial location for the fixation colour); 4. Complete mismatch (both the radial frequency pattern and spatial-location are incorrect relative to the fixation colour). The test display was shown for two seconds and then participants were asked separately about first the spatial-location match and then object match (both relative to the central fixation colour). Participants had 2 seconds to respond to each question and then they rated how confident they were about their response for both the spatial-location and object match using a slider (from 0-100%; untimed response). After this, they were presented with the correct pairing for 2 seconds, either confirming a correct response or providing a corrective learning opportunity for incorrect responses.

Task difficulty increased systematically across the four test blocks. In Block 1 (easiest), incorrect object trials differed from the correct radial frequency pattern by more than two points (e.g. a radial frequency with 5 vs 8 points), and incorrect location trials placed the stimulus in the orthogonal visual quadrant. In Blocks 2 and 3 (medium difficulty), incorrect objects differed by two radial frequency points, and incorrect locations were presented in one of the two quadrants adjacent to the correct location. In Block 4 (hardest), incorrect objects differed by only one radial frequency point, and incorrect locations were presented within the same quadrant, offset by 20° from the correct location (see Figure 2 and Supplementary Materials for example stimuli).

**Figure 2.**
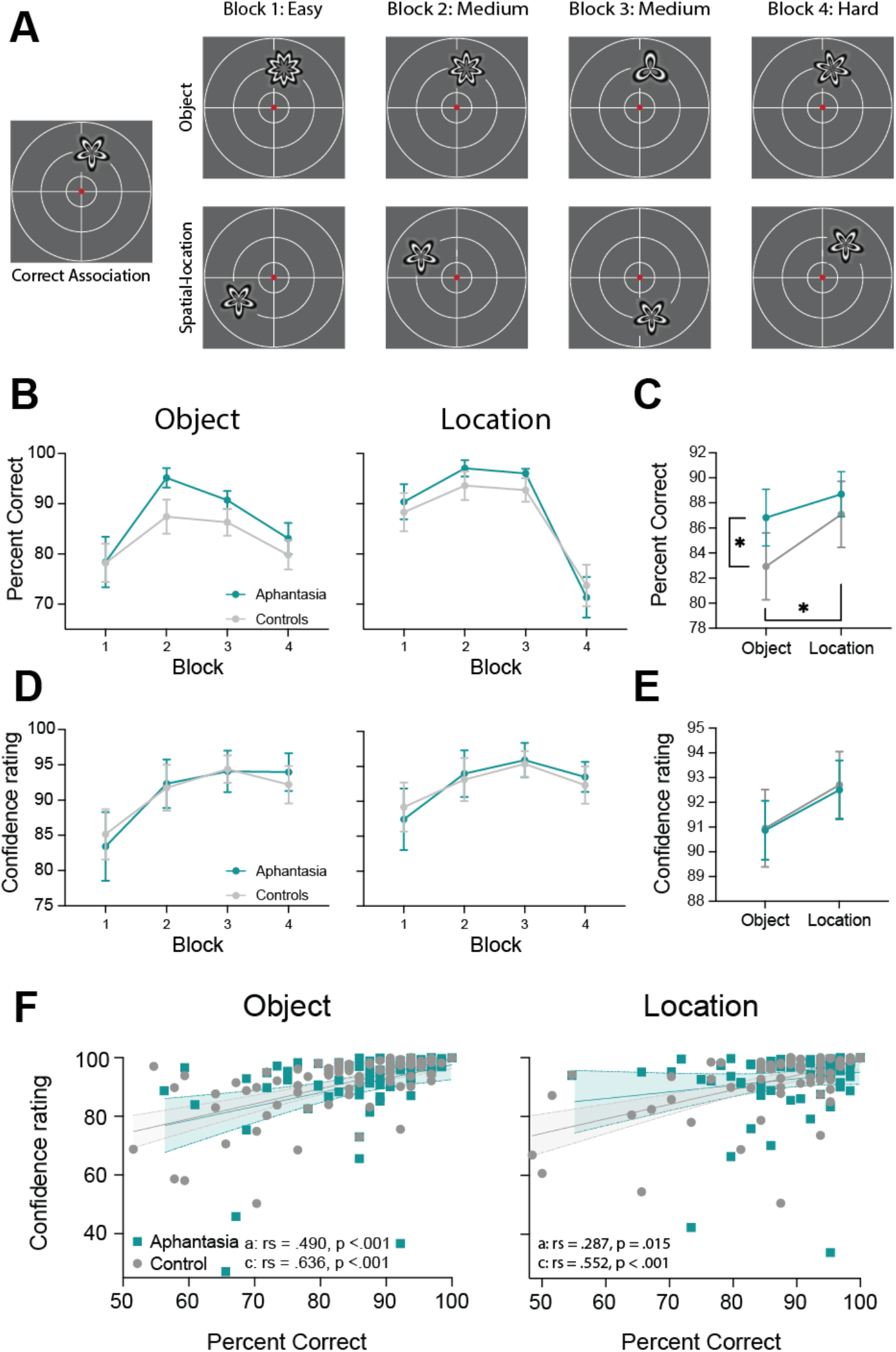
**A.** Schematic demonstrates the increasing difficulty of the task across the four blocks. Shows examples of incorrect object (top row) and incorrect spatial-location trials (bottom row) for each of the blocks/difficulty levels. **B**. Accuracy (% correct) for all 4 blocks for the object and location memory components for the aphantasia (teal) and control group (grey). **C**. Accuracy (% correct) averaged across the 4 blocks for the object and location memory components for the aphantasia (teal) and control group (grey). **D**. Confidence ratings for all 4 blocks for the object and location memory components for the aphantasia (teal) and control group (grey). **E**. Confidence ratings averaged across all 4 blocks for the object and location memory components for the aphantasia (teal) and control group (grey). **F**. Scatterplots show the relationship between confidence and performance for the object (left panel) and location (right panel) for the aphantasia group (teal) and control group (grey), averaged across all four blocks. For B-E means and 95% confidence intervals are presented.

### Post-test Strategy Questionnaires

After completing the full memory test (all 4 learning and 4 test blocks), participants were asked to select the strategy they used most frequently out of several options (see figure 2 and supplementary materials). They were then presented with three common strategies and asked to rate how often they used each of the strategies (5-point Likert scale) and how much they agreed with three statements regarding strategy switching and the ease of the task (5-point Likert scale, full wording of the questionnaire can be found in the supplementary materials).

## Results

### Pre-registered analysis

The accuracy data (see Figure 2 B & C for group data and Supplementary Figure S2 for individual subject data) were analysed with a 2 (Group: aphantasia & controls) x 2 (Memory type: object & spatial-location) x 4 (Block/difficulty: 1, 2, 3, 4) mixed repeated measures ANOVA. Age was added as a covariate for all analyses due to significant differences between the two groups (U = 260, *p* < .001) and Greenhouse-Geisser correction was applied where required due to violations of Sphericity. There was a main effect of Block (F(2.379, 336.187) = 6.80, *p* < .001, see Figure 2 B and Supplementary Table S1 for post-hoc block analysis) but no interaction between Block and Group (F(2.379, 336.187) = .581, p = .589).

There was a significant main effect of Group (F(1,146) = 7.33, *p* = .008) and an interaction between Group and Memory type (F(1,146) = 7.661, *p* = .006) when controlling for age (see Figure 2C for main effect averaged data). There was no significant three-way interaction (F(2.303, 336.187) = .638, *p* = .550).

Post-hoc analysis breaking down the group by memory type interaction showed that individuals with aphantasia were significantly better than controls for the object memory component of the task (*p* = .005, Bonferroni corrected) but not for the location task (*p* = .641, Bonferroni corrected, see Figure 2C). Post-hoc contrasts by group revealed that the control group was significantly better at spatial-location compared to object memory (*p* < .001, Bonferroni corrected), but this was not the case for the aphantasia group (*p* = .999, Bonferroni correction).

Given the importance of the lack of difference in our analyses above, and the significant age differences between the two groups, we conducted additional separate Bayesian repeated-measures ANOVAs for the aphantasia and control groups (not pre-registered). This allows us to assess the degree of evidence for either the alternative hypothesis of a difference (Bayes Factor (BF) > 1, with > 3 considered sufficient evidence of a difference for interpretation) or the null hypothesis of no difference (BF < 1, with <1/3 considered sufficient evidence for the null). In line with the frequentist omnibus analysis, there was weak evidence in the direction of a null effect when comparing object and spatial-location memory performance within the aphantasia group, BF_10_ = 0.67 (post hoc test), suggesting that memory performance did not differ reliably between conditions. In contrast, there was overwhelming evidence in favour of an effect of memory type in the control group, *BF*_10_ = 12,803 (post hoc test), showing that control participants performed better in the spatial-location condition compared to the object condition.

The confidence data were analysed with a 2 (Group: aphantasia & controls) x 2 (Memory Type: object & spatial-location) x 4 (block: 1, 2, 3, 4) mixed repeated measures ANOVA with age as a covariate (pre-registered analysis). There was no main effect of Group (F(1,146) = 1.691, *p* = .196) when controlling for age and no significant interactions between Group and any other factor (all *p* > .290; see Figure 2E for data averaged across block, for data for each block see Supplementary Figure 2). There was a significant interaction between Memory Type and Block with participants becoming more confident as the task went on (F(2.231, 325.667) = 4.290, *p* = .012, see supplementary table S2 for post-hoc comparisons).

We then assessed the degree to which confidence was associated with performance in each group by correlating the average performance for the object/spatial-location memory condition with the average confidence ratings for the same object/spatial-location condition. Due to violations of normality, Spearman’s rank correlations were performed (pre-registered correlations using frequentists statistics, Bayesian analysis was not pre-registered). For both groups, there were significant correlations between confidence and performance for the object (controls: r_s_ = .636, *p* < .001, tau = .464, BF_10_ = 6.26×10^6^; aphantasia: r_s_ = .490, *p* < .001, tau = .348, BF_10_ = 1513.2) and spatial-location memory conditions (controls: r_s_ = .552, *p* < .001, tau = .401, BF_10_ = 71561.82; aphantasia: r_s_ = .287, *p* = .015, tau = .195, BF_10_ = 2.782). Comparing the correlation coefficients, there were no significant differences between the two groups for either the object (t = -1.282, p = .199, BF_10_ = .187) or spatial-location correlations (t = -1.94, p = .052, BF_10_ = .540).

### Exploratory analysis of cognitive strategies

Due to our unexpected findings regarding the aphantasia group outperforming the control group for the object memory condition, we explored the reported cognitive strategies to assess whether the groups differed and whether strategy was associated with performance. We ran frequentist and Bayes’ analyses to measure the evidence for any effects. First, we assessed which strategy each group reported using most frequently (see Figure 3A). Control participants were most likely to endorse a visualising strategy (∼49%) as their most used strategy but almost as many reported using a verbal rehearsal strategy (∼42%; see Figure 3A). In contrast, individuals identifying as having aphantasia overwhelmingly reported using a verbal rehearsal strategy most frequently (∼90%).

**Figure 3.**
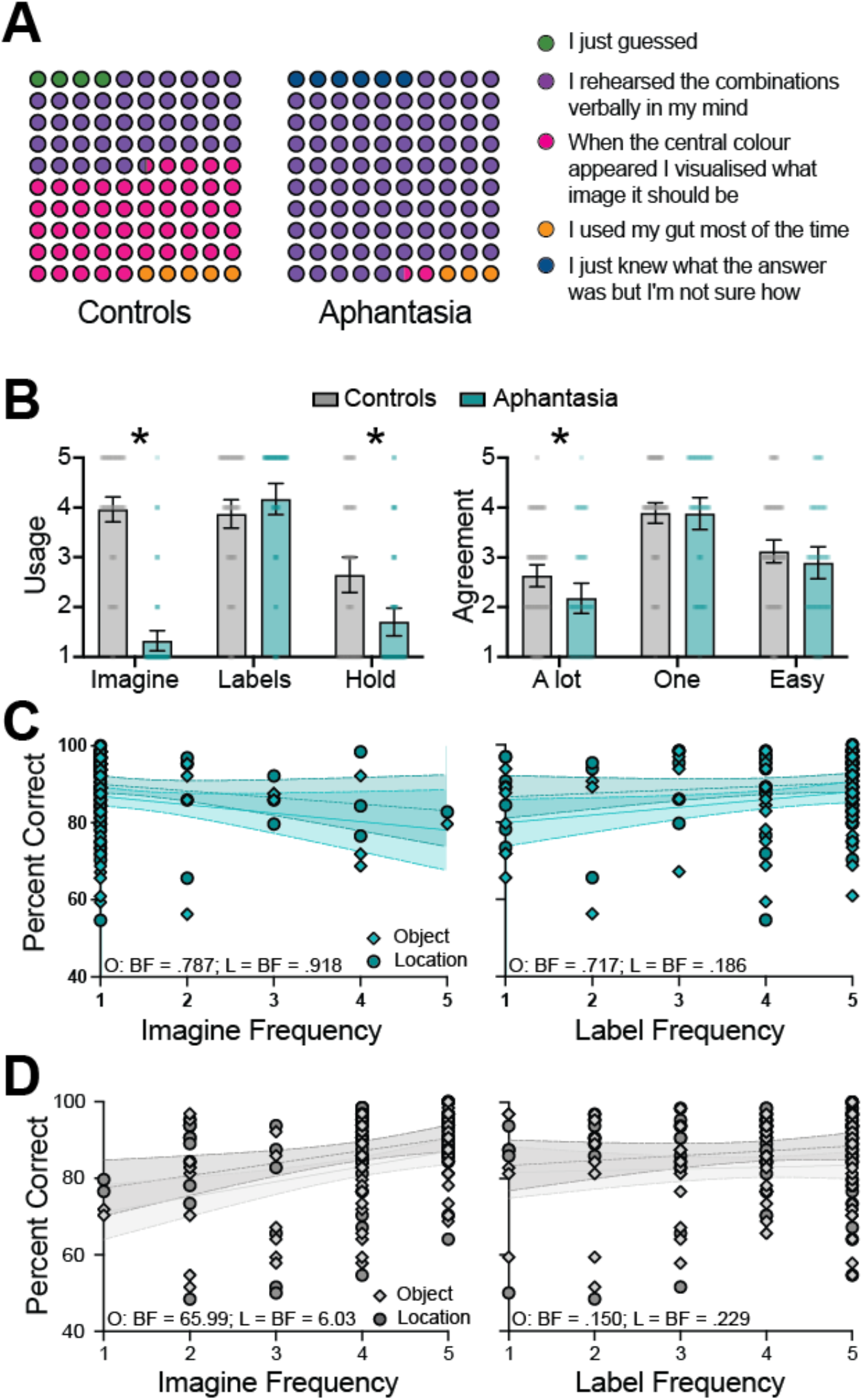
Strategy use in control and aphantasia groups. **A**. Strategy selected as best representing task performance by the control group (left panel) and the aphantasia group (right panel).**B**. Self-reported frequency of strategy use for *Imagine, Labels*, and *Hold* (left panel) questions (see questions below), and agreement with statements regarding strategy switching and task ease (right panel), shown for control (grey bars) and aphantasia (teal bars) groups.**C**. Scatterplots showing correlations between reported frequency of strategy use and performance in the aphantasia group across all four blocks: *Imagine* strategy use (left panel) and *Labels* strategy use (right panel) plotted against object (diamonds) and location (circles) performance. **D**. Scatterplots showing correlations between reported frequency of strategy use and performance in the control group across all four blocks: *Imagine* strategy use (left panel) and *Labels* strategy use (right panel) plotted against object (diamonds) and location (circles) performance. For all bars, means and 95% confidence intervals are shown. Asterisks indicate tests with *p* < .05 and Bayes factors > 3. O=object, L=location.

Participants were also asked 3 questions to assess the frequency with which they used visual imagery versus verbal labelling strategies (see Figure 3 and supplementary materials for wording of questions). Unsurprisingly, as a group, individuals with aphantasia were less likely to report using imagery-based strategies than the control group (Mann-Whitney t-tests, imagine question: W = 5172.5, *p* < .001, BF_10_ = 1.052×10^6^; hold question: W = 3656, *p* < .001, BF_10_ = 15.35, see Figure 3B and legend for questionnaire wording), and they were significantly more likely to report using a labelling type strategy (W = 2232, *p* = .037), although the Bayes Factor was inconclusive (BF_10_ = .509).

We also asked participants questions about strategy switching and ease of task and they rated how much they agreed with each statement on a 5-point Likert scale (see Figure 3 legend for wording of questions). Control participants were more likely to agree that they used a number of different strategies to help them remember the associations compared to the group with aphantasia (W = 3575, *p* < .001, BF_10_ = 10.91). There was no difference between the two groups regarding reporting sticking to one strategy (W = 2594, *p =* .572, BF_10_ = .209). Control participants were significantly more likely to agree that the task was easy compared to the aphantasia group (W = 3247.5, *p* = .027), but the Bayes factor suggested that the evidence for a group difference was inconclusive (BF_10_ = .963).

As the group with aphantasia performed better than controls and reported using nonvisual imagery-based strategies to perform the task, it could be the case that using nonvisual imagery strategies leads to better performance, rather than the group with aphantasia being better at the task *per se*. To assess this, we correlated each of the 6 strategies with performance, separately for the two groups, to see if there was evidence that any particular strategy correlated with better performance.

### Strategy items

#### Imagine

To remember the visual pattern and location I tried to imagine the image being at its specific location in my mind until the test image came up.

#### Labels

To remember the visual patterns, I rehearsed the images and locations of the images in my mind by giving them labels, e.g., I repeated green 2nd quadrant with a 3-point star

#### Hold

I tried to ‘hold’ the image on the correct location of the screen while I waited for the test image to come up.

#### A lot

I tried a lot of different strategies to remember the images and their locations. One: I chose one strategy and stuck to it to remember the images and their locations. Easy: I found this task very easy.

For the aphantasia group, none of the imagery or labelling strategy questions correlated with performance (all p > .122 and BF_10_ .186-.918, see Figure 3C). For the strategy switching questions, greater endorsement of using many strategies was negatively associated with object memory performance, r_s_ = -.271, p = .022, τ = -.214, BF_10_ = 4.808. Specifically, individuals with aphantasia who agreed that they used many strategies tended to perform worse on the object component of the task than those who disagreed. In line with the confidence ratings, there were significant positive correlations with those who reported finding the task easier performing better on both the object and spatial-location components of the task (object: r_s_ = .464, *p* < .001, tau = .195, BF_10_ = 3238.57; spatial-location: r_s_ = .476, *p* < .001, tau = .376, BF_10_ = 5130.24).

In the control group, participants who reported using an imagery strategy more frequently performed better on both the object and spatial-location conditions (object: r_s_ = .347, *p* = .002, tau = .273, BF_10_ = 65.991; spatial-location: r_s_ = .272, *p* = .017, tau = .213, BF_10_ = 6.033; see Figure 3C left panel). In contrast, reported use of a labelling strategy was not associated with performance for either memory type, with Bayes factors providing evidence in favour of the null (object: r_s_ = .012, *p* = .919, tau = .011, BF_10_ = .150; spatial-location: r_s_ = .092, *p* = .428, tau = .073, BF_10_ = .229, see Figure 3C right panel). The hold strategy also did not correlate significantly with performance in either condition (object: r_s_ =.092, *p* = .424, tau = .072, BF_10_ = .226; spatial-location: r_s_ = .174, *p* = .130, tau = .127, BF_10_ = .558). For the strategy switching questions neither question significantly correlated with performance (all *p* > .07 and BF_10_ .149-.1.43). Self-reported task ease was positively correlated with performance on both the object and spatial-location components of the task (object: r_s_ = .517, *p* < .001, tau = .411, BF_10_ = 146380.25; spatial-location: r_s_ = .435, *p* < .001, tau = .348, BF_10_ = 2933.70, respectively).

## Discussion

We investigated differences in associative visual memory for objects and spatial-locations between individuals who report lacking mental imagery (aphantasia) and those who report visual imagery (controls). We found that individuals with aphantasia performed significantly better than control participants on the object memory component of this task, contrary to predictions. There was no significant difference in spatial-location memory performance between the two groups. Further, individuals with aphantasia did not show a significant difference in their performance between the object and spatial-location memory conditions, with anecdotal evidence for a null effect. In contrast, controls performed better on the location memory component of the task compared to the object component. We found no significant differences in the mean confidence ratings between the two groups, contrary to our hypotheses. Both groups also showed good metacognitive insight with confidence ratings positively correlating with performance for both the object and spatial-location conditions.

These findings are surprising given previous work showing that individuals with aphantasia perform worse on memory for objects but no different on spatial information when drawing scenes from memory (Bainbridge et al., 2021), although a recent study also found no differences in performance for object versus location memory in aphantasia when using a more traditional continuous response task (Siena & Simons, 2024). One explanation for our finding of better object memory in aphantasia compared to controls may lie in the strategies employed by participants and our task design. Most individuals with aphantasia reported using nonvisual strategies, suggesting imagery is not required to perform this task. For the radial frequency stimuli used here, labelling features (e.g., counting and remembering the number of points) may be a more efficient strategy than forming a mental image. However, in the control group, participants who reported using imagery more frequently to remember the images performed better, with no such correlation found between performance and a labelling strategy. This suggests that when imagery is used in this task, it is effective for those who have visual imagery, but for those lacking imagery, nonvisual strategies may be more highly developed, producing a relative advantage here.

The conflicting findings regarding object and spatial memory in aphantasia (Bainbridge et al., 2021; Siena & Simons, 2024) might be due to the type of task used, tapping into different forms of memory which might require visual imagery. Our task was a two-alternative forced-choice paradigm which could rely on familiarity rather than explicit recall, whereas the Bainbridge et al., (2021) study relied heavily on recall where participants must draw from their memory. Although Siena and Simons (2024) also required recall in a method-of-adjustment task, their task provided a physical stimulus that participants could adjust to match their memory. It may be that differences in object versus spatial memory only arise when memory is purely generative (as in drawing tasks or when recalling personal memories), whereas when memory can be compared to an externally presented stimulus (as in method-of-adjustment paradigms or alternative-forced tasks) no such impairments in performance occur. At this point, it is not clear whether the inconsistency of our findings with these previous studies is due to differences in recall versus recognition memory, or task design differences.

Another possibility is that despite differences in reports, individuals with aphantasia are still using some form of unconscious imagery, analogous to blindsight (Michel et al., 2025), but see also for alternative views on unconscious imagery in aphantasia (McKilliam & Kirberg, 2025; Scholz et al., 2025). While it is possible that individuals with aphantasia have poor metacognitive insight into their cognitive strategies and are relying on unconscious imagery, our confidence results make this an unlikely explanation for our findings. Confidence ratings correlated with performance for both object and spatial components of the task in both the aphantasia and control groups, indicating insight into task performance. Instead, it seems more likely that individuals with aphantasia are using a different nonvisual strategy to remember the associations for both object and locations, and thus there is little difference in performance for these two components. Conversely, for participants using imagery, it may be that the location is more easily represented with visual imagery while the object requires more precision, creating a baseline difference in difficulty between the memory types. Additionally, our task always tested location first, suggesting that imagery may decay quicker than the non-imagery strategies used by individuals with aphantasia. This should not occur if the aphantasia group are unconsciously using imagery to guide their performance.

The selection of task used for this study was based on findings that remembering associations recruits low-level visual cortex during delay periods to represent memory locations (Favila et al., 2022). Specifically, Favila et al., (2022) found that when participants recalled the location of a given association, there was retinotopic BOLD activity aligned to the remembered location, including V1, with the activity spatially broader and lower amplitude during memory than during perception. This was taken as evidence that this task requires low-level sensory representation, which may rely on visual imagery. This raises the question as to what would be found in individuals with aphantasia. Despite their lack of subjective visual imagery, our group of individuals with aphantasia may still recruit early visual cortex during recall of the object location, consistent with findings from recent studies showing successful decoding of visual information in V1 even in the absence of reported imagery during imagery and memory tasks (Chang et al., 2025; Weber et al., 2024). However, these latter studies use neural decoding in which a machine learning algorithm is trained on the delay period (i.e., when imagining/remembering) and tested on the delay period, rather than cross-decoding from perception to imagery. For example, in the Chang et al. (2025) study, while they were able to decode the content of colour imagery for individuals with aphantasia from V1 when training on imagery, they were unable to cross-decode the content of imagery when training a classifier on perception. This suggests that while the early visual cortex might be recruited during these tasks, this recruitment is not ‘perception-like’ enough to be decodable from perceiving these stimuli.

Another possibility is that the location of the stimuli might be represented retinotopically in individuals with aphantasia due to their preserved spatial imagery, but the object features themselves are stored in a nonvisual format. This would presumably recruit retinotopic cortex for location representations and other nonvisual regions for object representations. Alternatively, successful performance on our task for individuals with aphantasia might be due to recruitment of completely different neural regions reflecting their different (non-imagery) strategies, potentially involving spatial, semantic or language-related regions. Functional neuroimaging studies could help disentangle these possibilities, shedding light on the flexibility and diversity of neural mechanisms supporting memory and imagery across individuals.

These findings have broader implications for understanding the impact of having aphantasia. They suggest that aphantasia may reflect differences in memory strategies rather than a fundamental impairment, and that individuals with aphantasia can maintain accurate metacognitive insight into their performance. While our task shows intact associative memory for simple, highly labelable stimuli, it remains to be seen whether these findings generalise to more complex or dynamic memory stimuli. Future work combining behavioural testing with neuroimaging could clarify which neural circuits support object and spatial memory in aphantasia. Further, our findings indicate that self-reported aphantasia does not impair the ability to form associative memories. This suggests that previously reported autobiographical memory deficits in aphantasia (Dawes et al., 2020; Dawes et al., 2022; Greenberg & Knowlton, 2014; Milton et al., 2021; Zeman et al., 2015) are unlikely to be driven by a general inability to form associations. Stimuli that are more difficult to label may still reveal deficits, but general associative memory appears intact.

Several limitations to our study should be noted. First, as with any special population study motivation may have differed between groups. Participants in the aphantasia group self-selected due to interest in the topic, while the control group participated as a course requirement. However, performance for location memory was equivalent, and age differences (controls were younger) should favour the control group. Second, we relied on self-report to classify aphantasia. The VVIQ was used to measure the lack of imagery in aphantasia here and, while commonly used, it was not designed for the assessment of aphantasia; better tools are needed for future studies. We also used an online study, which allows much larger samples but has limitations regarding testing conditions, however our rigorous pre-registered exclusions meant that we ensured participants who were not completing the task properly were removed from the dataset. Lastly, while we did present several different frequently reported cognitive strategies for participants to rate their use of, we did not provide an open-ended question about strategies and thus we might have missed some of the alternative strategies individuals used to remember the images that were not verbal or visual in nature.

Overall, this study provides clear evidence that self-reported deficits in visual imagery do not result in impairments in associative memory. In fact, individuals reporting aphantasia seem to be, if anything, better able to remember object associations than those reporting typical imagery. This study also provides evidence that individuals with aphantasia retain good metacognitive insight into their memory performance suggesting that aphantasia is not just a case of generally poor metacognition or poor introspection. Taken together, these results suggest that visual imagery is not necessary for effective associative memory, at least of stimuli that can be labelled. Other non-imagery-based strategies may support successful encoding and retrieval when imagery is unavailable, highlighting the often-hidden heterogeneity of human mental experiences.

## CRediT Author Statement

**RK:** Conceptualization, Methodology, Investigation, Formal analysis, Visualization, Writing – Original Draft, Funding acquisition.

**ZI:** Conceptualization, Writing – Review & Editing.

**ANR:** Conceptualization, Methodology, Supervision, Writing – Review & Editing.

## Acknowledgments

We would like to thank participants with aphantasia for taking the time to participate in these studies. The funding for this project was from an MQRF to RK MQRF0001061-2020; RK is supported by an ARC DECRA DE240100606; ANR is supported by an ARC FT230100119.

