## Supplementary Materials for "Associative Visual Memory in Aphantasia: Evidence for Intact Object and Spatial Memory, Metacognitive Awareness, but Different Strategies"

Cognitive strategy Questions:

Please choose the following strategy you used the most to help you perform the task

- I just guessed
- I rehearsed the combinations verbally in my mind
- When the central colour appeared, I visualised what image it should be
- I used my gut most of the time
- I just knew the answer but I'm not sure how

Imagine: To remember the visual pattern and location I tried to imagine the image being at it's specific location in my mind until the test image came up

1 = never used this strategy 2 3 4 5 = used it all the time

Label: To remember the visual patterns I rehearsed the images and locations of the images in my mind by giving them labels, i.e. I repeated green 2nd quadrant with a 3-point star

1 = never used this strategy 2 3 4 5 = used it all the time

'Hold': I tried to 'hold' the image on the correct location of the screen while I waited for the test image to come up

1 = never used this strategy 2 3 4 5 = used it all the time

A lot: I tried a lot of different strategies to remember the images and their locations

1 = totally disagree 2 3 4 5 = totally agree

One: I chose one strategy and stuck to it to remember the images and their locations

1 = totally disagree 2 3 4 5 = totally agree

Easy: I found this task very easy

1 = totally disagree 2 3 4 5 = totally agree

**All Stimuli Used**

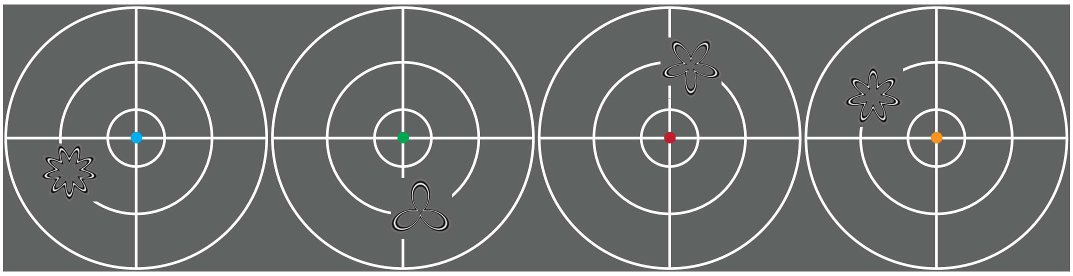

Learnt Associations

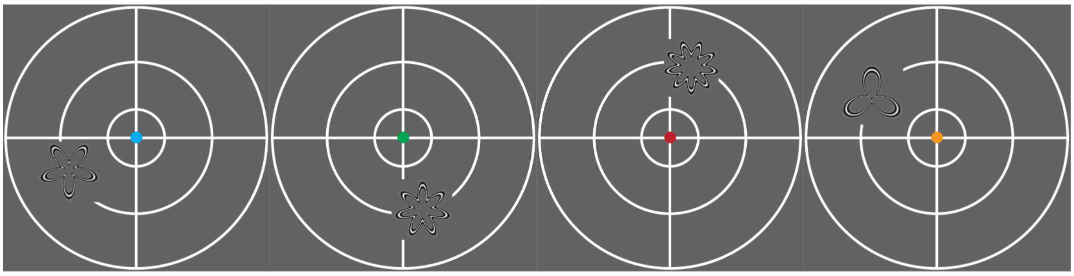

Correct Location Block 1

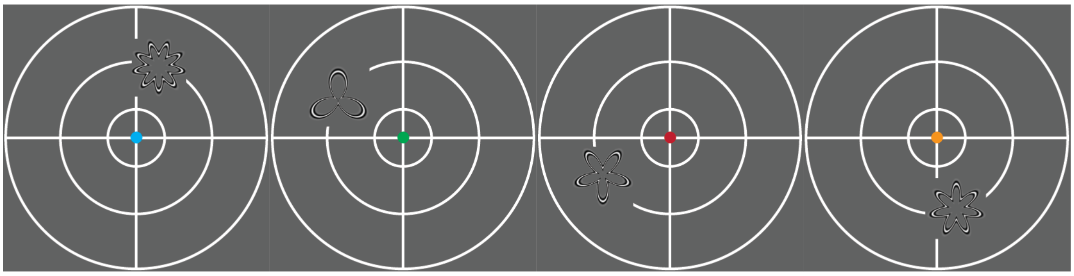

Correct Pattern Block 1

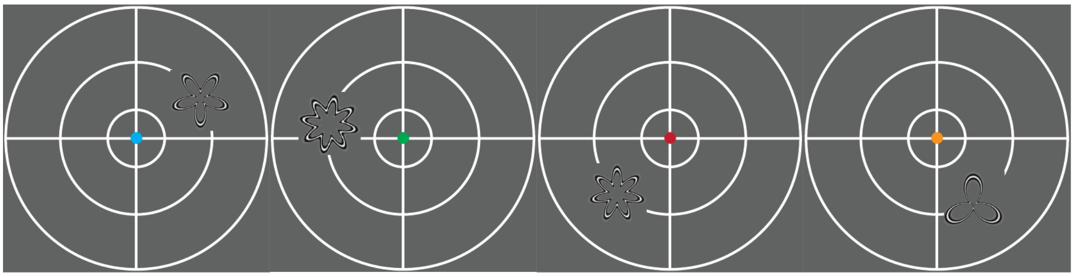

Incorrect all Block 1

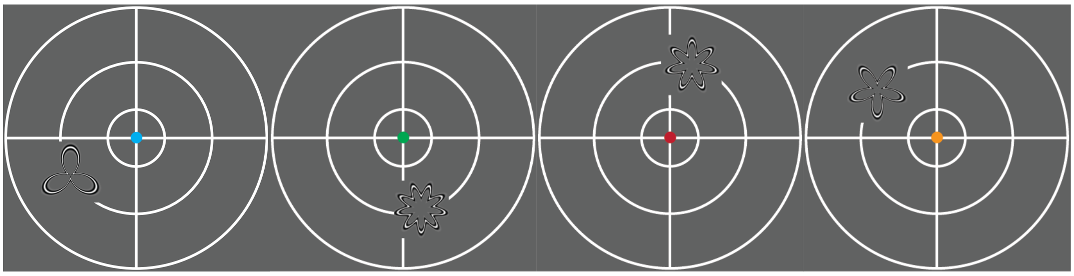

Correct Location Block 2

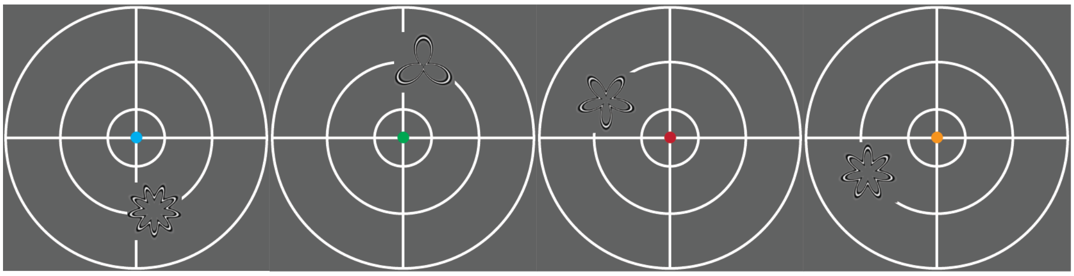
Correct Pattern Block 2

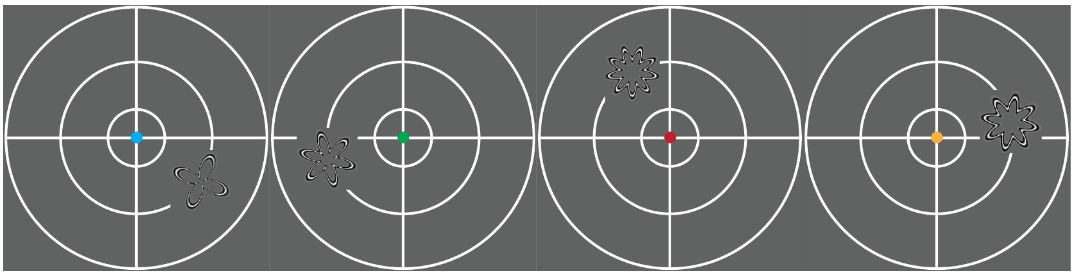
Incorrect all Block 2

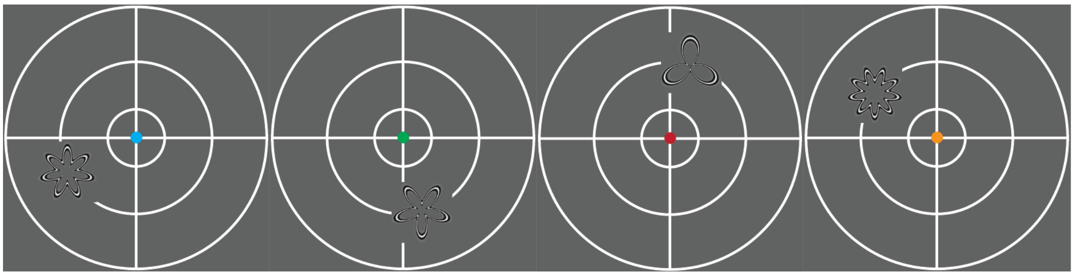
Correct Location Block 3

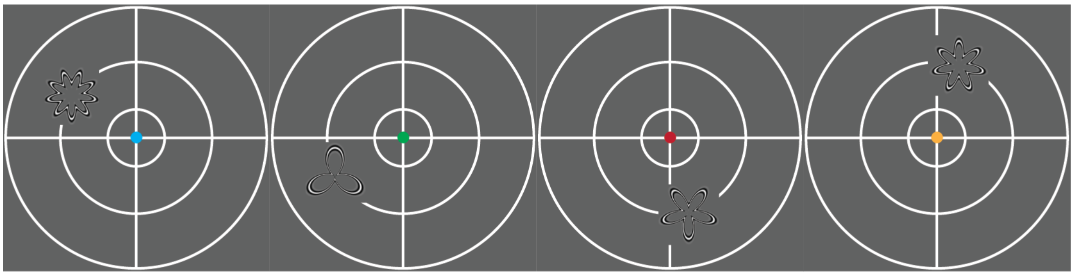
Correct Pattern Block 3

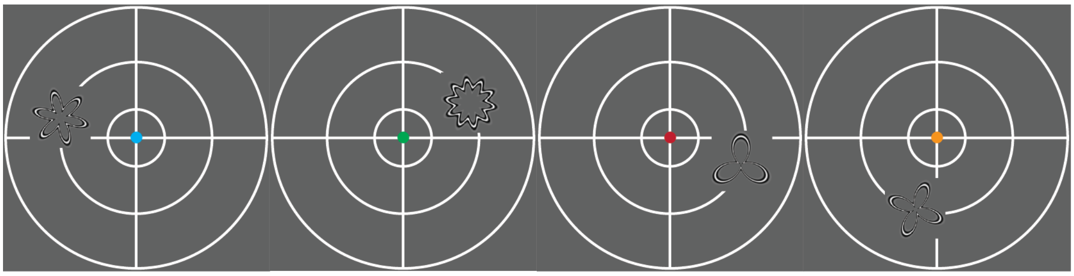
Incorrect all Block 3

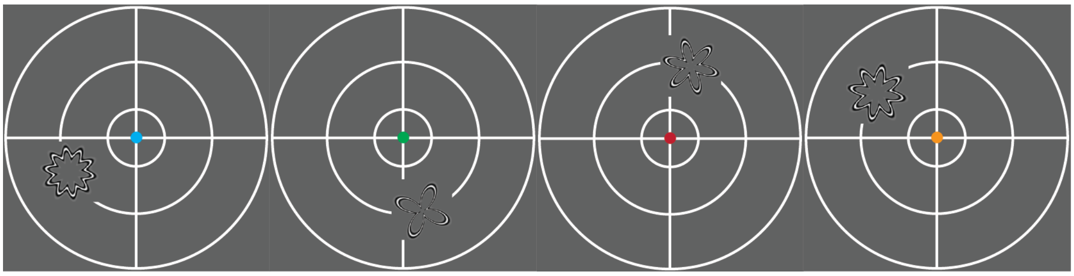

Correct Location Block 4

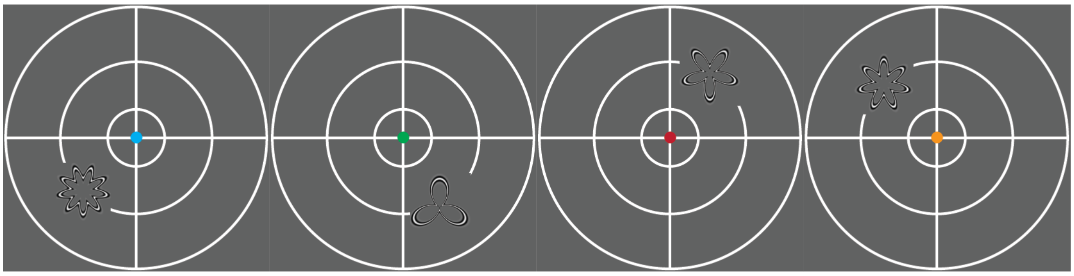

Correct Pattern Block 4

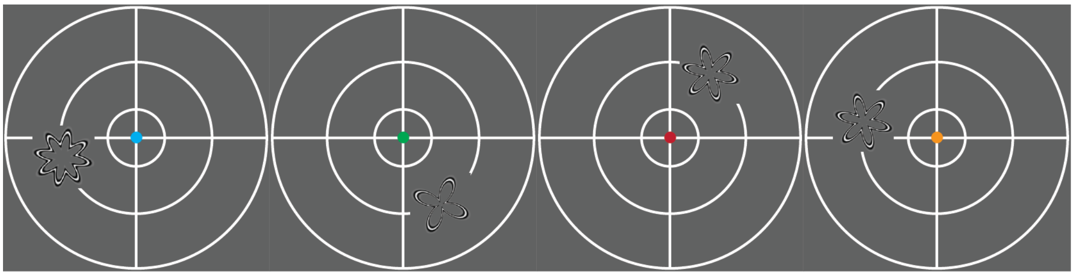

Incorrect all Block 4

### Supplementary Figures

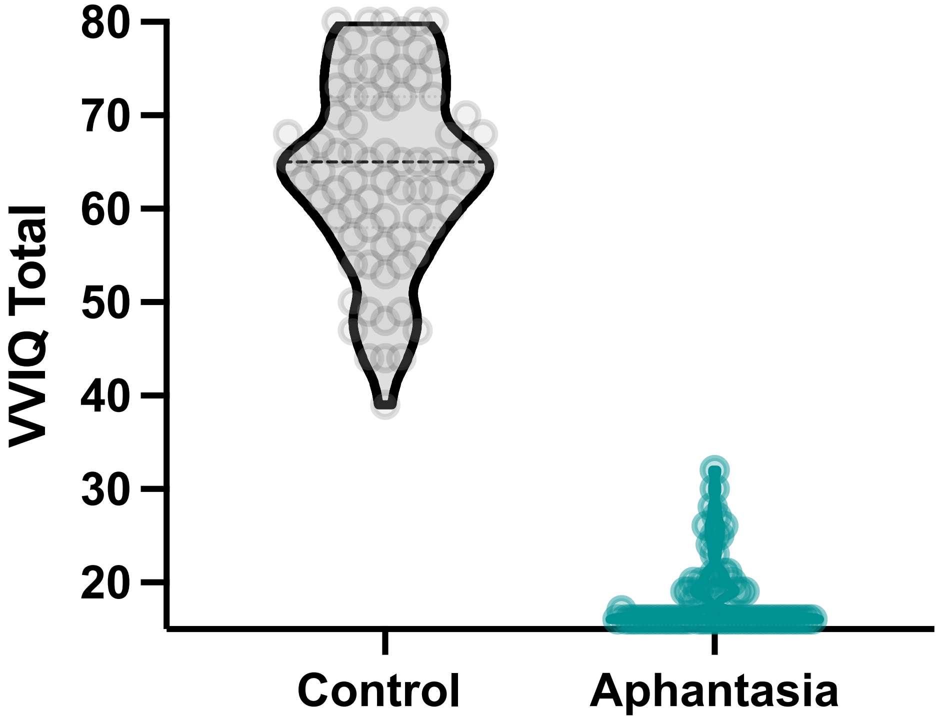


**Supplementary Figure S1**. VVIQ scores for the control (grey) and aphantasia (teal) groups.

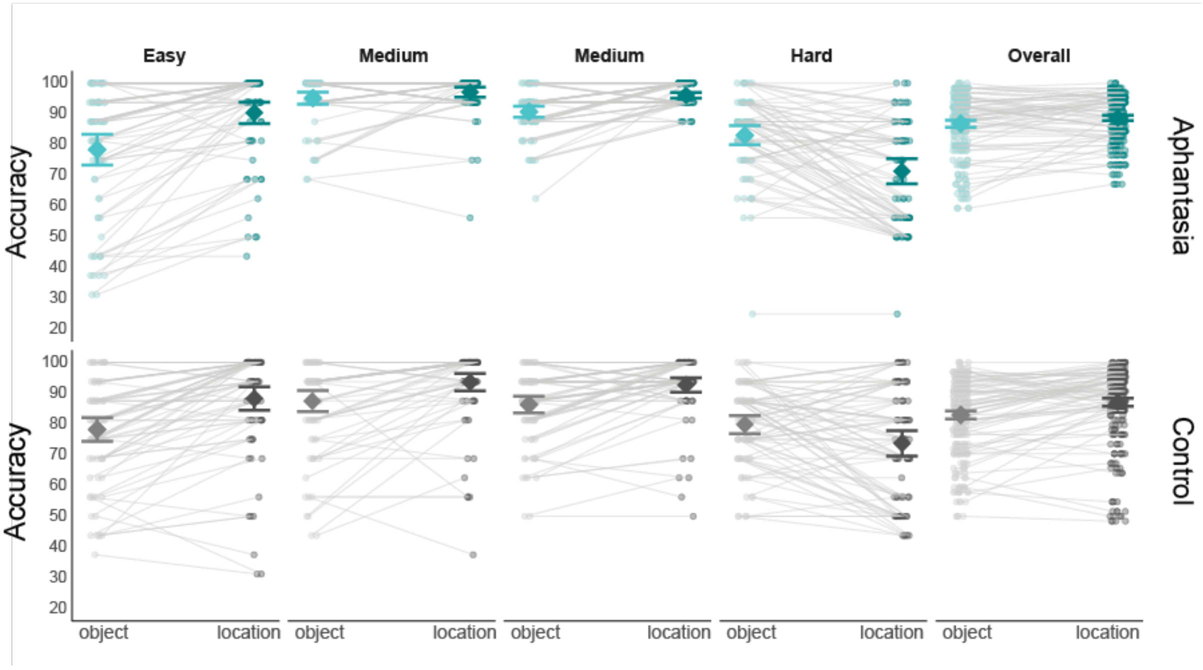


**Supplementary Figure S2**. Performance scores for the aphantasia (teal) and control (grey) groups per block/difficulty level. Individual data points represent each participant's score, and grey lines connect the same participant across conditions. Large symbols represent the group mean, with 95% confidence intervals.

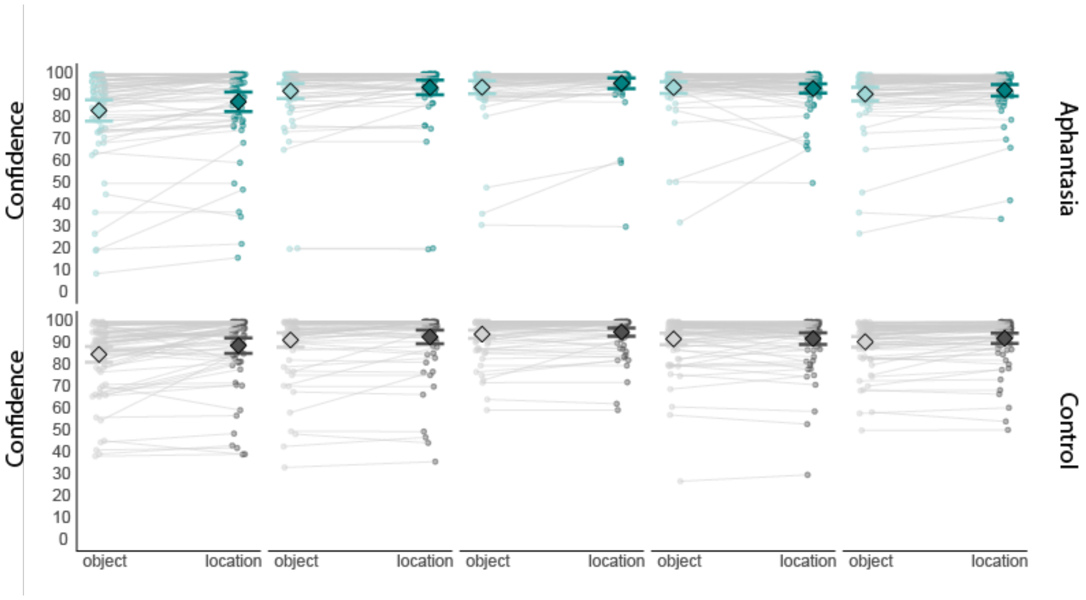


**Supplementary Figure S3**. Confidence scores for the aphantasia (teal) and control (grey) groups per block/difficulty level. Individual data points represent each participant's score, and grey lines connect the same participant across conditions. Large symbols represent the group mean, with 95% confidence intervals.

### Supplementary Tables

**Supplementary Table S1:** Table shows the post-hoc tests for performance as a function of Block, following up the main effect of Block.

*Post Hoc Comparisons - Block*

| | | Mean Difference | SE | t | $p_{bonf}$ |
|---|---|---|---|---|---|
| 1 | 2 | -9.555 | 1.007 | -9.487 | < .001 |
| | 3 | -7.704 | 1.105 | -6.969 | < .001 |
| | 4 | 6.865 | 1.269 | 5.412 | < .001 |
| 2 | 3 | 1.851 | 0.655 | 2.825 | 0.032 |
| | 4 | 16.420 | 1.071 | 15.326 | < .001 |
| 3 | 4 | 14.569 | 0.915 | 15.923 | < .001 |

*Note.* P-value adjusted for comparing a family of 6
*Note.* Results are averaged over the levels of: Group, Type

**Supplementary Table S2:** Table shows the post-hoc tests for the interaction between Block and Memory type for confidence.

*Post Hoc Comparisons - Memory Type ✻ Block*

| | | Mean Difference | SE | t | $p_{bonf}$ |
|---|---|---|---|---|---|
| Location, 1 | Object, 1 | 4.012 | 0.595 | 6.743 | < .001 |
| | Location, 2 | -5.175 | 1.053 | -4.916 | < .001 |
| | Object, 2 | -3.671 | 1.057 | -3.472 | 0.019 |
| | Location, 3 | -7.237 | 1.084 | -6.675 | < .001 |
| | Object, 3 | -5.844 | 1.076 | -5.431 | < .001 |
| | Location, 4 | -4.507 | 1.289 | -3.497 | 0.017 |
| | Object, 4 | -4.711 | 1.219 | -3.865 | 0.005 |
| Object, 1 | Location, 2 | -9.187 | 1.256 | -7.312 | < .001 |
| | Object, 2 | -7.683 | 1.177 | -6.526 | < .001 |
| | Location, 3 | -11.248 | 1.219 | -9.229 | < .001 |
| | Object, 3 | -9.856 | 1.206 | -8.171 | < .001 |
| | Location, 4 | -8.518 | 1.440 | -5.914 | < .001 |
| | Object, 4 | -8.722 | 1.340 | -6.511 | < .001 |
| ***Location, 2*** | Object, 2 | 1.504 | 0.340 | 4.419 | < .001 |
| | ***Location, 3*** | ***-2.061*** | ***0.707*** | ***-2.914*** | ***0.116*** |
| | ***Object, 3*** | ***-0.669*** | ***0.723*** | ***-0.925*** | ***1.000*** |
| | ***Location, 4*** | ***0.669*** | ***0.810*** | ***0.826*** | ***1.000*** |
| | ***Object, 4*** | ***0.465*** | ***0.779*** | ***0.597*** | ***1.000*** |
| **Object, 2** | Location, 3 | -3.566 | 0.743 | -4.800 | < .001 |
| | ***Object, 3*** | ***-2.173*** | ***0.701*** | ***-3.100*** | ***0.065*** |
| | ***Location, 4*** | ***-0.835*** | ***0.823*** | ***-1.016*** | ***1.000*** |
| | ***Object, 4*** | ***-1.040*** | ***0.762*** | ***-1.365*** | ***1.000*** |
| Location, 3 | Object, 3 | 1.392 | 0.323 | 4.313 | < .001 |
| | Location, 4 | 2.730 | 0.645 | 4.230 | 0.001 |
| | Object, 4 | 2.526 | 0.622 | 4.060 | 0.002 |
| ***Object, 3*** | ***Location, 4*** | ***1.338*** | ***0.680*** | ***1.968*** | ***1.000*** |

*Post Hoc Comparisons - Memory Type ✲ Block*

| | | Mean Difference | SE | t | $p_{bonf}$ |
|---|---|---|---|---|---|
| | ***Object, 4*** | ***1.134*** | ***0.582*** | ***1.949*** | ***1.000*** |
| ***Location, 4*** | ***Object, 4*** | ***-0.204*** | ***0.465*** | ***-0.439*** | ***1.000*** |

*Note.* P-value adjusted for comparing a family of 28
*Note.* Results are averaged over the levels of: Group
